# Targeting the Kidney’s Osmotic Gradient with AVPR2 Antagonism Limits Inflammation and Fibrosis after Injury

**DOI:** 10.64898/2026.10.21.760644

**Authors:** Karen I. López-Cayuqueo, Felix J. Boivin, Shuang Cao, Janna Leiz, Larissa Nassauer, Axel Schambach, Christian Hinze, Kai M. Schmidt-Ott

**Author notes:** **Corresponding author:** Prof. Dr. med. Kai Schmidt-Ott, Department of Nephrology and Hypertension, Hannover Medical School (MHH), 30625 Hannover, Germany.

## Abstract

**Background:** The renal medulla is characterized by a steep osmotic gradient that is essential for urinary concentration but may also influence cellular stress and injury responses. While vasopressin-dependent medullary hyperosmolarity is well established in water homeostasis, its role in shaping kidney injury and repair remains poorly understood.

**Methods:** We pharmacologically reduced vasopressin signaling using the Avpr2 antagonist Tolvaptan and examined the consequences for renal medullary osmolality, tubular stress responses, inflammation, and fibrotic remodeling. Single-nucleus RNA sequencing, bulk transcriptomics, quantitative PCR, and histological analyses were performed in healthy kidneys and following unilateral ischemia-reperfusion injury (IRI) or cisplatin-induced nephropathy.

**Results:** Tolvaptan treatment lowered renal medullary osmolality and suppressed osmotic stress-associated transcriptional programs in medullary collecting duct cells. Following IRI, Tolvaptan-treated mice exhibited attenuated expression of inflammatory and monocyte recruitment genes, reduced macrophage accumulation in the renal medulla, and diminished fibrotic remodeling at later stages. In cisplatin-induced nephropathy, Tolvaptan reduced tubular injury markers and inflammatory activation in the outer stripe of the outer medulla.

**Conclusions:** These findings identify vasopressin-dependent renal medullary hyperosmolarity as a modulator of tubular stress, inflammation, and fibrotic remodeling after kidney injury. Modulation of the medullary osmotic environment influences early injury trajectories and downstream repair responses across distinct models of acute kidney injury.

## INTRODUCTION

Acute kidney injury (AKI) remains a major clinical problem and is increasingly recognized as a driver of chronic kidney disease (CKD) through maladaptive repair and fibrotic remodeling^1,2^. Although substantial progress has been made in defining inflammatory and epithelial pathways involved in kidney injury, the contribution of the kidney’s unique physiological microenvironment to these processes is less well understood.

One of the most distinctive features of the kidney is the steep corticomedullary osmotic gradient, which enables urinary concentration. In the renal medulla, interstitial osmolality far exceeds that of other tissues and is dynamically regulated by vasopressin signaling through the arginine vasopressin receptor 2 (Avpr2) in collecting duct principal cells^3^. This hyperosmolar environment imposes substantial adaptive demands on medullary epithelial cells and is associated with the activation of osmoprotective and stress-related transcriptional programs^4,5^.

Beyond its role in water balance, medullary hyperosmolarity has been implicated in shaping immune cell behavior. Experimental studies have demonstrated that elevated sodium and osmolality can modulate mononuclear phagocyte recruitment, polarization, and antimicrobial function within the kidney medulla^6,7^. These observations raise the possibility that the osmotic environment of the renal medulla is not merely permissive but actively participates in injury and repair processes.

Whether vasopressin-dependent medullary hyperosmolarity influences the kidney’s response to injury remains unclear. In particular, it is unknown whether modulation of medullary osmolality alters tubular stress responses, inflammatory activation, and the progression from acute injury to fibrotic remodeling. Addressing this question requires an integrated analysis of renal physiology, cell-type-specific transcriptional responses, and tissue-level injury outcomes.

Here, we used pharmacological Avpr2 antagonism with Tolvaptan to reduce renal medullary osmolality and examined its impact on tubular stress programs, inflammation, and fibrosis.

Using single-nucleus RNA sequencing and complementary molecular and histological approaches, we assessed injury responses in both ischemia-reperfusion injury and cisplatin-induced nephropathy. Our findings indicate that vasopressin-dependent medullary hyperosmolarity shapes injury-associated inflammatory and remodeling responses across distinct models of AKI.

## METHODS

### Mice

Adult male C57BL/6N mice were used for all experiments. Mice were housed under specific pathogen-free conditions on a 12-hour light–dark cycle with *ad libitum* access to food and water. Tolvaptan-treated animals received chow containing 0.1% (w/w) Tolvaptan (LKT Laboratories, T5847; CAS 150683-30-0), homogeneously mixed into standard rodent diet, as previously described^8^. Control animals (hereafter referred to as placebo-treated) received identical chow without the drug. Tolvaptan treatment was initiated 48 hours prior to experimental procedures and maintained throughout the study period. All animal experiments were approved by the responsible authorities (LAGeSo Berlin and LAVES Hannover, Germany) and were conducted in accordance with institutional and EU guidelines for animal care and use.

### Urine osmolality measurements

Spot urine samples were collected immediately before euthanasia and stored at −80°C until analysis. Urine osmolality was determined by freezing-point depression using a freezing-point osmometer (Gonotec, Germany).

### Tissue osmolality measurements

Renal tissue osmolality was determined as previously described^9^, with minor modifications. Kidney regions were microdissected immediately after harvest, and wet tissue weight was recorded. Samples were subsequently dried under vacuum overnight to determine dry weight. Lyophilized tissue was re-weighed and then boiled in 200 μl ultrapure H₂O for 1 hour, followed by centrifugation at 13,000 × g for 10 minutes. The supernatant was collected and used for osmolality measurement by freezing-point depression. Tissue osmolality was calculated according to the formula: Tissue osmolality = (osmolality of supernatant x volume of water added)/ (wet weight−dry weight). Results are expressed as milliosmoles per kilogram of H₂O (mOsm/kg H₂O).

### Ischemia-reperfusion injury

Unilateral renal ischemia–reperfusion injury (IRI) was performed in male C57BL/6N mice (8–12 weeks of age). Mice were anesthetized with isoflurane (2.3% in air; flow rate 350 ml/min) and placed on a temperature-controlled heating pad (Physitemp Instruments, Clifton, NJ, USA) to maintain body temperature at 37°C. Core temperature was continuously monitored using a rectal probe.

Following midline abdominal incision, the left renal pedicle was carefully isolated and occluded using a nontraumatic microaneurysm clip (FE690K, Aesculap, Berlin, Germany) for 25 minutes. Successful ischemia and subsequent reperfusion were confirmed visually. After clip removal, the incision was closed in layers. Postoperatively, mice received analgesia with buprenorphine (0.1 mg/kg, subcutaneous).

Animals were sacrificed at either 7 or 21 days after reperfusion. Both the ischemic and contralateral kidneys were harvested for molecular and histological analyses.

### Cisplatin-induced nephropathy

For induction of toxic acute kidney injury, mice received cisplatin (9 mg/kg body weight; intraperitoneal injection) or an equivalent volume of vehicle once weekly for three consecutive weeks. Tolvaptan-treated mice remained on Tolvaptan-containing chow throughout the experimental period, beginning 48 hours prior to the first cisplatin injection. Control mice received identical cisplatin injections while maintained on placebo chow.

Animals were sacrificed 72 hours after the third cisplatin injection. Blood was collected at sacrifice for measurement of serum creatinine, and urea. Kidneys were harvested for molecular analyses and histological assessment, including quantification of tubular injury markers and inflammatory cell infiltration.

### Creatinine and urea concentrations

Whole blood samples were allowed to clot at room temperature for 20 min, followed by centrifugation at 8,000 rpm for 10 min (Centrifuge 5425 R, Eppendorf). The resulting serum was collected, and creatinine and urea concentrations were measured using an enzymatic assay in an AU480 clinical chemistry analyzer (Beckman Coulter) at the Animal Phenotyping Facility of the Max Delbrück Center (MDC, Berlin, Germany).

### Generation of single-nucleus RNA sequencing libraries

Immediately after sacrifice, kidneys were harvested and placed in ice-cold PBS. A 1-2 mm transverse slice containing cortex, outer medulla, and inner medulla was dissected as previously described^10,11^. Tissue samples were incubated in RNAlater (Invitrogen, AM7020) at 4°C for 24 hours and subsequently stored at -80°C until processing.

For nuclei isolation, frozen tissue was minced in ice-cold nuclear lysis buffer (Sigma) supplemented with 1 U/μl RiboLock RNase inhibitor and 10 mM ribonucleoside vanadyl complex (RVC). Tissue was homogenized using a Dounce homogenizer (Sigma D8938– 1SET) with pestle A, passed through a 100 μm filter, further homogenized with pestle B, and filtered through a 40 μm cell strainer. The homogenate was centrifuged at 500 × g for 5 minutes at 4°C.

The nuclear pellet was resuspended in nuclear lysis buffer supplemented with 1 U/μl RiboLock. To remove debris, the suspension was underlaid with lysis buffer containing 10% sucrose and 1 U/μl RiboLock and centrifuged at 500 × g for 5 minutes at 4°C. The supernatant and debris layer were carefully removed. Purified nuclei were resuspended in PBS containing 0.04% BSA and 1 U/μl RiboLock, filtered through a 20 μm strainer, and stained with 4’,6-diamidino-2-phenylindole (DAPI).

Nuclei were sorted using a BD FACSAria II cell sorter to exclude debris, damaged nuclei, and multiplets. Sorted nuclei were counted and processed for single-nucleus RNA sequencing using the Chromium Single Cell 3′ Reagent Kit v3.1 (10x Genomics), targeting 10,000 nuclei per sample. Libraries were sequenced on a NovaSeq 6000 platform (Illumina) using paired-end sequencing. Digital gene expression matrices were generated using Cell Ranger (v3.0.2) with the parameter --force-cells=10000.

### Single-nuclei sequencing data processing and clustering

Single-nucleus RNA sequencing datasets (placebo-treated, n = 2; Tolvaptan-treated, n = 2) were processed in R^12^ using Seurat^13^ (v4.0.4). Raw count matrices (10x Genomics output) were imported using Read10X and converted into Seurat objects (CreateSeuratObject; min.features = 500; min.cells = 3). Mitochondrial transcript content was calculated using PercentageFeatureSet (pattern “^mt-”). Nuclei were retained if they contained 600-8,000 detected genes and <2.5% mitochondrial transcripts. After filtering, datasets were merged and split by sample identity (orig.ident). Each sample was log-normalized (NormalizeData; LogNormalize; scale factor 10,000) and variable features were identified (FindVariableFeatures; “vst”; 2,000 features).

Samples were integrated using anchor-based integration (FindIntegrationAnchors and IntegrateData; dims 1–30). Unless otherwise indicated, clustering and visualization were performed on the integrated assay. Data were scaled (ScaleData), principal component analysis was performed (RunPCA; 30 PCs), and UMAP was computed using the first 30 principal components (RunUMAP; dims 1–30). A shared nearest-neighbor graph was constructed (FindNeighbors; dims 1–30) and clustering was performed using FindClusters (resolution = 0.5).

For cell-type annotation, marker genes conserved across samples were identified using FindConservedMarkers (grouping.var = “orig.ident”; min.pct = 0.25; logfc.threshold = 0.25). Iterative subclustering was performed for selected cell populations by subsetting the integrated object and repeating neighbor graph construction and clustering, followed by manual annotation based on established markers.

Differential expression between Tolvaptan-and placebo-treated nuclei was assessed using FindMarkers on the RNA assay (min.pct = 0.05; logfc.threshold = 0.58). Data visualization and figure preparation were performed using Seurat and additional R packages including ggplot2, dplyr, dittoSeq, and patchwork.

### Bulk RNA sequencing and data processing

Bulk RNA sequencing was performed on whole kidneys obtained from adult male C57BL/6N mice (placebo-treated, n = 5; Tolvaptan-treated, n = 6). Tolvaptan treatment was initiated 48 hours prior to unilateral ischemia-reperfusion injury (IRI), and kidneys were collected 7 days after reperfusion.

Total RNA was extracted and processed according to the service provider’s protocols (Novogene). Libraries were prepared for mRNA sequencing and sequenced using paired-end 150 bp reads on an Illumina NovaSeq 6000 platform.

Raw FASTQ files were aligned to the mouse reference genome (mm9) using STAR aligner. Gene-level counts were generated using featureCounts with exon-level summarization (-p -t exon -g gene_id -s 0). For visualization, gene expression values were converted to transcripts per million (TPM), log2-transformed as log2(TPM + 1), and subsequently rescaled to a range of 0-1 using min-max normalization independently for each gene across samples.

Differential gene expression analysis was performed using DESeq2^14^ (v1.40.2) in R on bulk RNA-sequencing count data. Gene expression was compared between Tolvaptan-and Placebo-treated mice, with log2 fold changes calculated as Tolvaptan relative to Placebo. Genes with an adjusted p-value < 0.05 were considered differentially expressed, without applying an additional fold-change magnitude cutoff. Differential expression results were visualized using volcano plots generated with ggplot2^15^ (v4.0.3). Gene Ontology (GO) biological process over-representation analysis was performed using Enrichr^16,17^ through the enrichR (v3.4) R package (GO Biological Process 2025). Genes with significantly lower expression in Tolvaptan-treated kidneys (adjusted p < 0.05; log2FC < 0) were included in the enrichment analysis. All genes with valid DESeq2 adjusted p-values (n = 14,997) were used as the background gene set. GO terms with enrichment-adjusted p-values < 0.05 were considered significant. Selected representative enriched biological processes were visualized as -log10(adjusted enrichment p-value), with the number of contributing genes indicated for each term. For human kidneys, publicly available bulk transcriptomic datasets^18^ from individuals with acute kidney injury (AKI) compared with non-AKI controls were used (adjusted p < 0.05, fold change ≤ 0.5).

### RNA *in situ* hybridization

Fluorescent in situ hybridization was performed using the RNAscope Multiplex Fluorescent Reagent Kit v2 (Advanced Cell Diagnostics [ACD], Newark, CA; Cat. No. 323110) according to the manufacturer’s instructions. Kidneys were fixed in 4% paraformaldehyde in PBS for 24 hours at room temperature with gentle agitation, dehydrated through a graded ethanol series, and embedded in paraffin. Paraffin-embedded tissue was sectioned at 5 μm thickness and mounted on Superfrost™ Plus slides. Sections were air-dried overnight, baked at 60°C for 1 hour, cooled to room temperature, deparaffinized, and air-dried prior to assay initiation. Target-specific probes were used for the following transcripts: Cryab (ACD #552741), Ddit3 (ACD #317661), Hmgb1 (ACD #406691), Nupr1 (ACD #434811), and Aqp2 (ACD #452411-C3). Pretreatment, hybridization, amplification, and detection steps were performed according to the RNAscope Multiplex Fluorescent Reagent Kit v2 protocol with sample preparation and pretreatment guidelines (ACD document #323100-USM). All probes were processed under identical conditions. Sections were counterstained with DAPI and mounted using Fluorescence Mounting Medium (Dako, S3023). Images were acquired using a ZEISS PALM MicroBeam microscope.

### Masson’s trichrome staining and fibrosis quantification

Following harvest, kidneys were sectioned into 1-mm transverse slices and fixed in 4% paraformaldehyde for 1 hour on ice. Samples were processed for paraffin embedding according to standard protocols. Paraffin sections (5 μm) were deparaffinized and rehydrated through graded alcohols. Masson’s trichrome staining was performed using a commercial Trichrome Stain kit (Sigma-Aldrich) according to the manufacturer’s instructions. Sections were mounted using Eukitt mounting medium (Sigma-Aldrich). Tubulointerstitial fibrosis was quantified by measuring the percentage of collagen-positive area in 10–12 non-overlapping fields per section at 200× magnification. Image acquisition was performed using a Leica DM2000 LED bright-field microscope. Quantification was conducted in a blinded manner using ImageJ^19^ with identical threshold settings applied across all samples.

### Immunofluorescence staining and quantification

Paraffin-embedded kidney sections (2 μm) were deparaffinized and rehydrated through graded alcohols. Antigen retrieval was performed in pre-warmed 10 mM citrate buffer pH 6 (>98°C) for 15 minutes, followed by cooling at room temperature for 30 minutes. Sections were blocked for 2 hours at room temperature in 5% normal donkey serum containing 0.25% Triton X-100 in PBS. Primary antibodies were diluted in 1% BSA in PBS and incubated overnight at 4°C. After primary incubation, sections were washed three times for 10 minutes each in PBS containing 0.25% Triton X-100. Secondary antibodies were applied for 2 hours at room temperature. For identification of proximal tubules and delineation of cortical regions, sections were co-stained with FITC-conjugated Lotus tetragonolobus lectin (LTL; 1:200; Vector Laboratories, FL-1321-2) diluted in PBS and incubated for 1 hour at room temperature. Nuclei were counterstained with DAPI (1:1000; Invitrogen, D1306). Sections were mounted using Dako Fluorescence Mounting Medium (Dako, S3023). For quantitative analysis, 10–12 non-overlapping fields per section were acquired using identical imaging settings across all experimental groups. Fluorescence-positive area was quantified in a blinded manner using Fiji^20^, applying identical threshold parameters to all samples. Confocal images were acquired using a Zeiss LSM980 laser scanning confocal microscope.

A complete list of antibodies and reagents is provided in Supplementary Table S1.

### RNA isolation, cDNA synthesis, and quantitative RT-PCR

Total RNA was isolated from kidney tissue using the NucleoSpin RNA Midi Kit (Macherey-Nagel, Cat. No. 740962.20) according to the manufacturer’s instructions. RNA concentration and purity were assessed using a DeNovix DS-7 spectrophotometer (DeNovix).

For cDNA synthesis, 1 µg of total RNA was reverse-transcribed using random primers (Promega, C118A), 5× RT buffer (Promega, M531A), dNTPs (Promega, U151A), and M-MLV reverse transcriptase (Promega, M170B), following the manufacturer’s protocol.

For quantitative real-time PCR (qRT-PCR), cDNA was diluted 1:10 in nuclease-free water. Amplification was performed using FastStart Taq DNA Polymerase (Roche, 04738420001) and SYBR Green I dye (Invitrogen, S7563) on a LightCycler® 96 system (Roche). Cycling conditions consisted of an initial denaturation step at 95°C for 10 minutes, followed by 45 cycles of 95°C, 60°C, and 72°C (three-step amplification protocol).

Specificity of amplification was verified by melt curve analysis and agarose gel electrophoresis of PCR products. Relative gene expression was calculated using the ΔΔCt method and normalized to hypoxanthine-guanine phosphoribosyltransferase (Hprt) as the housekeeping gene.

Primer sequences are listed in Supplementary Table S2.

### Tumor cell viability assays

To investigate whether tolvaptan affects cisplatin-induced cytotoxicity in tumor cells, *in vitro* experiments were performed using three human cancer cell lines obtained from the American Type Culture Collection (ATCC) and subsequently modified to express mCherry: CaSki and SiHa (cervical carcinoma) and OVCAR3 (ovarian carcinoma). Cells were treated with cisplatin (10 μM) in the presence or absence of Tolvaptan (1 μM or 3 μM) for 72 hours. Cell proliferation was assessed over time by live-cell imaging using the CELLCYTE X™ (Cytena). The tumor cell lines were seeded at a density of 2x10^4^ cells per well in 96-well plates and treated with Tolvaptan and cisplatin the following day. Images were captured from each condition every 2 hours using a 4x objective in the red and enhanced-contour channel. Flow cytometric analyses were performed to quantify mCherry-positive tumor cells, indicating viable cell populations, and additionally, to evaluate apoptotic/dead cells via Annexin V assay. For this, 5x10^4^ cells were seeded in 12-well plates and treated with cisplatin or Tolvaptan 24 hours after. For the flow cytometric analysis, dead cells from the medium, as well as adherent cells, were harvested and stained with FITC-Annexin V (BD Biosciences, Cat. No. 560931) in Annexin V Binding Buffer (BD Biosciences) for 10 minutes at room temperature in the dark. Subsequently, cells were stained with DAPI and analyzed in the BD FACSCanto™ flow cytometer.

### Statistics

All RNA-seq data were analyzed using R^12^, while all other data were processed with GraphPad Prism 10. Statistical tests, along with sample sizes (n), are specified in the corresponding figure legends. A P value of less than 0.05 was considered significant.

## RESULTS

### Tolvaptan reduces renal medullary osmolality and suppresses osmotic stress-associated transcriptional programs

To determine how pharmacological inhibition of vasopressin signaling affects the renal medullary environment, adult male C57BL/6N mice were treated with the AVPR2 antagonist Tolvaptan (1 mg/g chow) for 48 hours prior to analysis (Figure 1A). As expected from effective blockade of vasopressin-dependent water reabsorption, Tolvaptan treatment reduced urine osmolality (Figure 1B). Consistent with this physiological change, Tolvaptan significantly reduced renal medullary osmolality, whereas cortical osmolality remained largely unchanged (Figure 1C).

**Figure 1.**
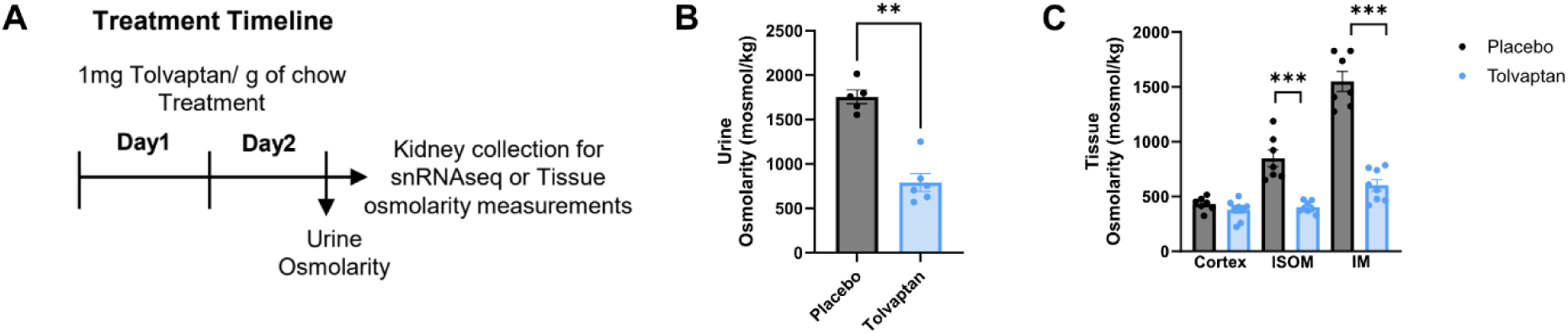
Effects of Tolvaptan on urine and renal tissue osmolality. **(A)** Experimental design: adult male C57BL/6N mice were treated with the AVPR2 antagonist Tolvaptan (1 mg/g chow) for 48 hours prior to analysis or Placebo (control diet without Tolvaptan). **(B)** Urine osmolality measured in Placebo-and Tolvaptan-treated mice. **(C)** Tissue osmolality measured in kidney cortex, inner stripe of the outer medulla (ISOM), and inner medulla (IM) from Placebo-and Tolvaptan-treated mice. Each dot represents an individual mouse. Data are shown as mean ± SEM, statistical significance was assessed using a Mann-Whitney U test comparing Tolvaptan versus placebo in each kidney region (**P < 0.01, ***P < 0.001).

To characterize cell type-specific transcriptional responses to Tolvaptan treatment, single-nucleus RNA sequencing was performed on kidneys from Tolvaptan-and Placebo-treated mice. Quality control metrics indicated comparable data quality and sequencing depth across samples (Supplementary Fig. S1). Unsupervised clustering followed by refinement identified all major renal epithelial, endothelial, and immune cell populations (Figure 2A). Cell identities were assigned based on established marker gene expression (Figure 2B). The relative abundance of major cell types and refined subclusters was comparable between Tolvaptan-and Placebo-treated groups, indicating that short-term AVPR2 inhibition did not substantially alter overall renal cellular composition (Figure 2B).

**Figure 2.**
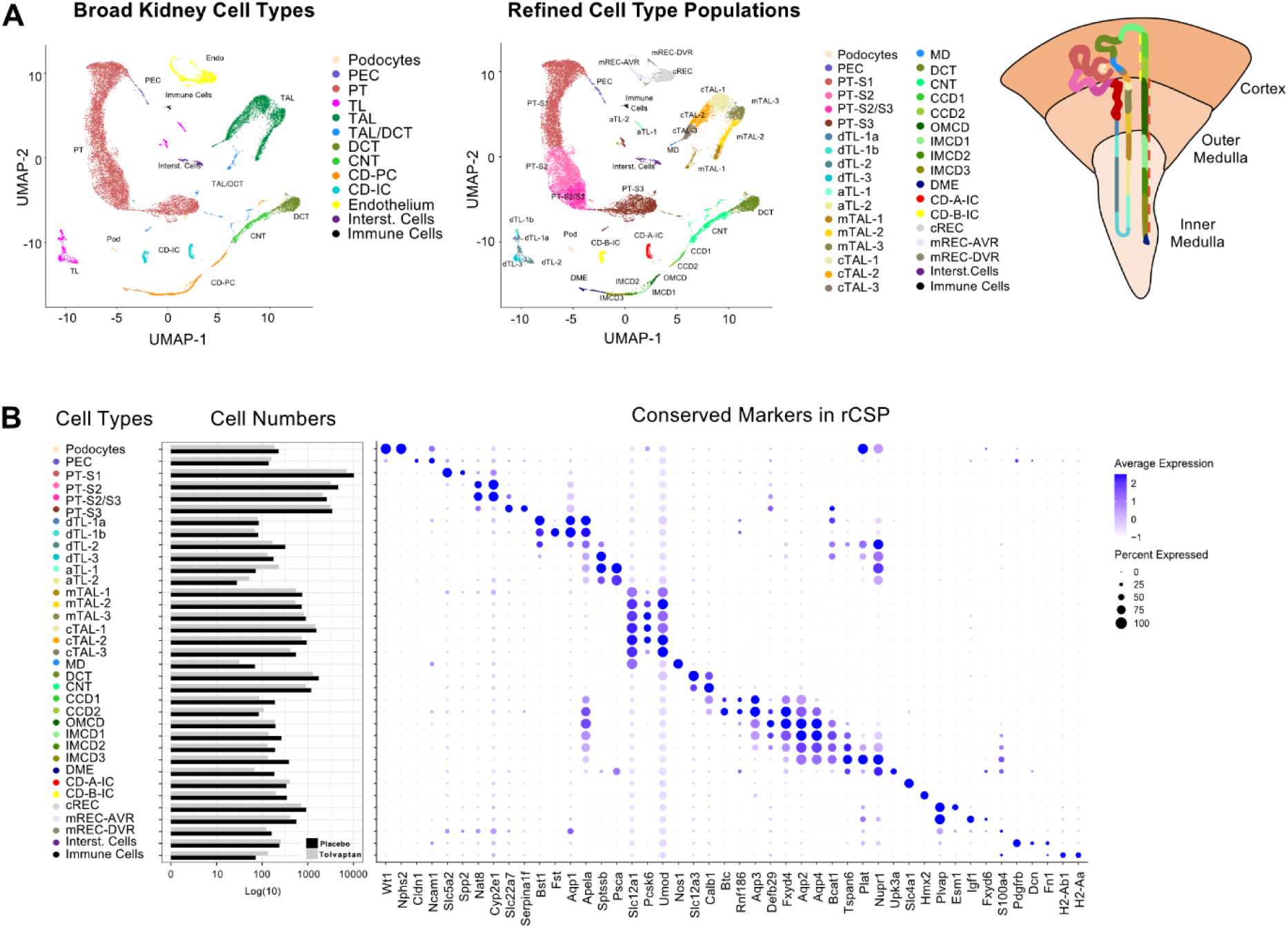
Single-nucleus RNA sequencing of mouse kidney identifies major cell types and refined subpopulations following Tolvaptan treatment. **(A)** Unbiased identification of kidney cell types. Uniform manifold approximation and projection (UMAP) visualization of integrated single-nucleus RNA sequencing data from pooled kidney nuclei across all experimental samples (n=2 Tolvaptan-treated and n=2 Placebo-treated mice). Each point represents a single nucleus transcriptome, and colors indicate cluster identity. Thirteen major renal cell types were identified based on established marker genes, followed by annotation of 35 refined cellular subpopulations (rCSPs). A schematic representation of the nephron illustrates the localization of rCSPs. **(B)** Quantification of the relative abundance of major renal cell types and refined subclusters (right), alongside a dot plot showing expression of established marker genes used for cell type annotation (left). Dot size represents the fraction of cells expressing each gene, and color intensity indicates average expression level. **Abbreviations:** aTL1/2, ascending thin limb cluster 1 and 2; CCD1/2, cortical collecting duct cluster 1 and 2; CD-A/B-IC, collecting duct intercalated cells Alpha/Beta; CD-PC, collecting duct principal cells; CNT, connecting tubule; cREC, cortical renal endothelial cells; cTAL1/2/3, cortical thick ascending limb clusters 1-3; DCT, distal convoluted tubule; DME, deep medullary epithelium; dTL1/2/3, descending thin limb clusters 1-3; IMCD1/2/3, inner medullary collecting duct clusters 1-3; MD, macula densa; mREC-AVR/DVR, medullary renal endothelial cells (ascending/descending vasa recta); mTAL1/2/3, medullary thick ascending limb clusters 1-3; OMCD, outer medullary collecting duct; PEC, parietal epithelial cell; PT, proximal tubule; PT-S1/2/3, proximal tubule segments 1-3; TAL, thick ascending limb; TL, thin limb.

Within collecting duct populations, Tolvaptan treatment resulted in a pronounced reduction in the expression of established vasopressin-and osmolarity-responsive genes. Expression of *Aqp2*, a canonical downstream effector of AVPR2 signaling, was significantly reduced across medullary collecting duct subclusters, including outer medullary collecting duct (OMCD) and inner medullary collecting duct (IMCD) principal cells. This was accompanied by decreased expression of *Cryab*, a molecular chaperone previously linked to tissue hyperosmolarity^5^ (Figure 3A). *In situ* hybridization was performed on kidney sections, confirming reduced expression of *Aqp2* and *Cryab* (Figure 3B).

**Figure 3.**
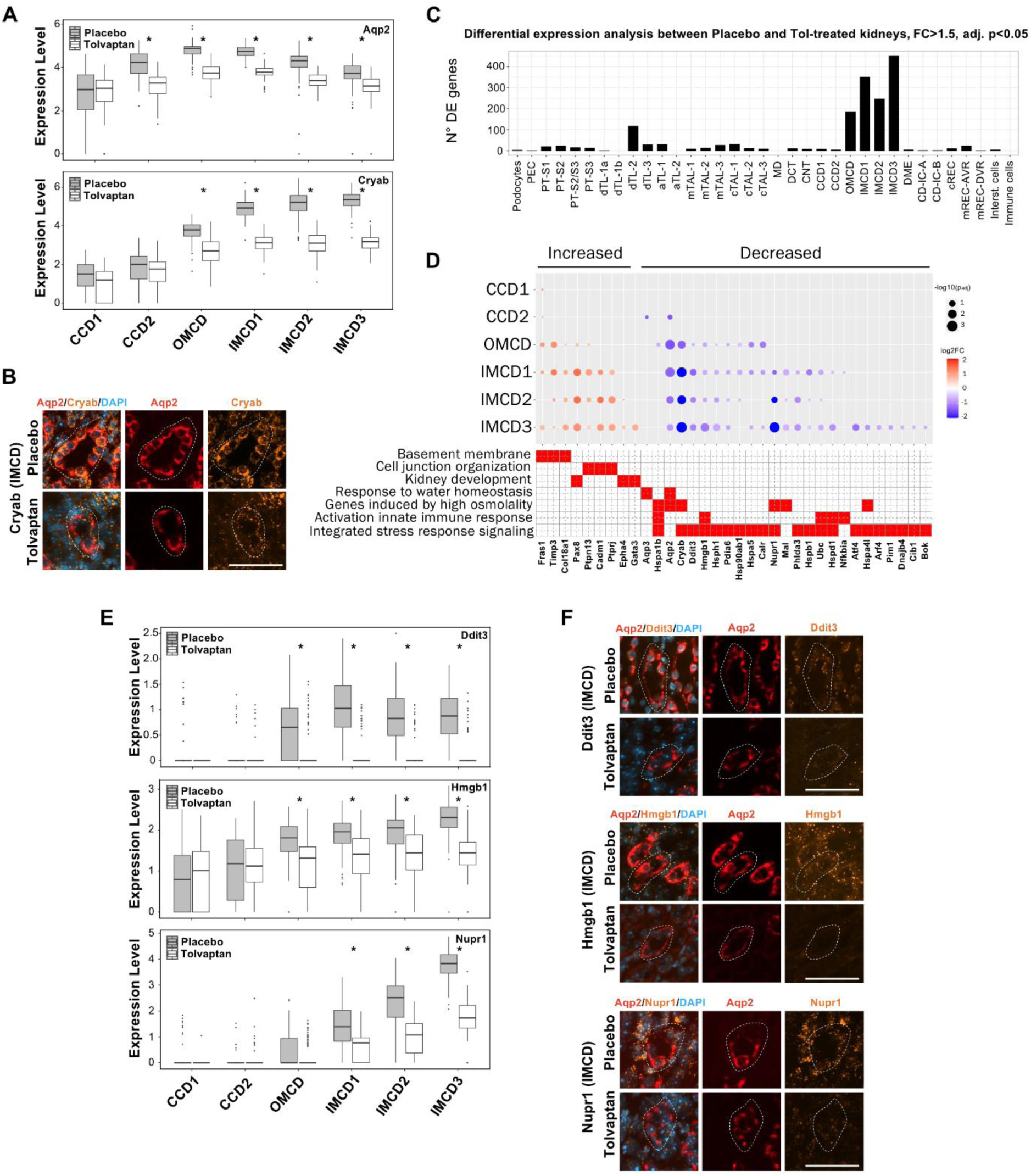
Transcriptional effects of Tolvaptan on osmotic stress-associated programs in collecting duct cells. **(A)** Expression of vasopressin-and osmolarity-responsive genes (*Aqp2, Cryab*) across collecting duct subclusters, including cortical collecting duct (CCD), outer medullary collecting duct (OMCD), and inner medullary collecting duct (IMCD), in Placebo-and Tolvaptan-treated mice. **(B)** Representative *in situ* hybridization images of *Aqp2* and *Cryab* expression in inner medullary collecting duct (IMCD) epithelia from Placebo-and Tolvaptan-treated mice (n = 3 mice per condition; scale bar, 50 µm). **(C)** Distribution of differentially expressed (DE) genes across nephron segments in Placebo-and Tolvaptan-treated mice. DE genes were defined by a fold change >1.5 and an adjusted *P* value < 0.05 (*), Bonferroni correction. **(D)** Gene Ontology enrichment analysis of genes downregulated in Tolvaptan-treated mice. Dot size represents the adjusted *P* value, and color indicates fold change. **(E)** Expression of stress-associated genes (*Nupr1, Ddit3, Hmgb1*) across collecting duct subclusters (CCD, OMCD, IMCD) in Placebo-and Tolvaptan-treated mice. **(F)** Representative *in situ* hybridization images of *Ddit3, Hmgb1,* and *Nupr1* expression in IMCD epithelia from Placebo-and Tolvaptan-treated mice (n = 3 mice per condition; scale bar, 50 µm).

Beyond classical vasopressin and osmolarity-responsive genes, differential expression analysis across refined subclusters revealed that the majority of significantly regulated genes were localized to medullary collecting duct compartments, with minimal transcriptional changes observed in cortical segments (Figure 3C). Gene Ontology enrichment analysis of downregulated transcripts identified significant overrepresentation of pathways related to hyperosmotic stress responses, activation of innate immune signaling, and integrated stress response pathways (Figure 3D).

Tolvaptan treatment was associated with reduced expression of several stress-associated genes, including *Nupr1*, *Ddit3*, and *Hmgb1*, predominantly within OMCD and IMCD populations (Figure 3E). To validate these findings spatially, *in situ* hybridization was performed on kidney sections, confirming reduced expression of *Ddit3*, *Hmgb1*, and *Nupr1* within medullary collecting duct epithelia of Tolvaptan-treated mice (Figure 3F).

To examine whether genes responsive to Tolvaptan in the mouse kidney are also associated with human kidney injury, we analyzed published kidney transcriptomic data from individuals with severe AKI and non-AKI controls^18^. Expression of *CRYAB*, *DDIT3*, and *NUPR1*, which was reduced following Tolvaptan treatment in mouse medullary collecting duct cells, was higher in human AKI samples (Supplementary Fig. S2A). Genes upregulated in human AKI were also enriched for Gene Ontology terms related to cellular stress and inflammatory responses (Supplementary Fig. S2B). These observations identify an overlap in selected stress-associated transcripts between the murine response to Tolvaptan and human AKI.

Together, these data demonstrate that Tolvaptan treatment not only attenuates canonical AVPR2 signaling but broadly suppresses osmotic stress-associated transcriptional programs in medullary collecting duct cells, and that similar programs are induced in human acute kidney injury.

To facilitate interrogation of the complete single-nucleus RNA sequencing dataset, we provide an interactive web-based resource that allows gene-by-gene visualization of expression changes across all refined renal cell populations following Tolvaptan treatment (see Data Availability).

### Reduction of medullary osmolality attenuates inflammatory responses after ischemia-reperfusion injury

To investigate whether modulation of renal medullary osmolality influences injury-associated inflammatory responses, we subjected Tolvaptan-and Placebo-treated mice to unilateral renal ischemia-reperfusion injury (IRI) and analyzed kidneys during the early recovery phase (7 days post-injury) (Figure 4A). This time point was selected to capture inflammatory and reparative processes preceding overt fibrotic remodeling.

**Figure 4.**
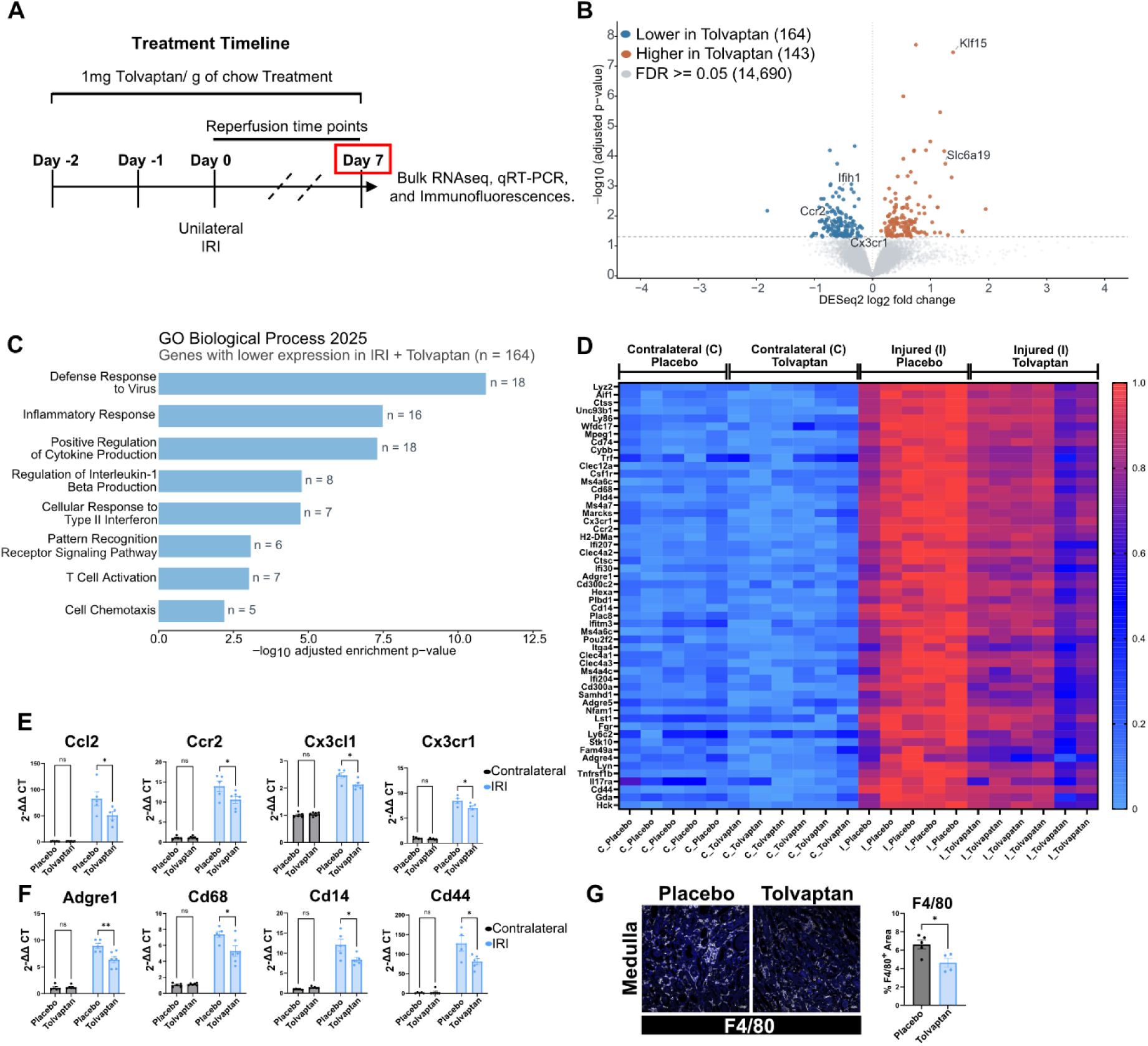
Effects of Tolvaptan treatment on the renal inflammatory response following ischemia-reperfusion injury. **(A)** Experimental design. Adult male C57BL/6N mice received Tolvaptan (1 mg/g chow) or Placebo (control diet without Tolvaptan) starting 48 hours before unilateral renal ischemia-reperfusion injury (IRI). Kidneys were collected 7 days after reperfusion for bulk RNA sequencing, quantitative real-time PCR (qRT-PCR), and immunofluorescence analyses. **(B)** Volcano plot showing differential gene expression in IRI kidneys collected from Tolvaptan-and Placebo-treated mice. Differential expression was assessed using DESeq2. The x-axis represents log2 fold change (Tolvaptan versus Placebo), and the y-axis shows −log10(adjusted p-value). Genes with significantly lower (blue) or higher (orange) expression in Tolvaptan-treated kidneys are indicated (adjusted p < 0.05), while nonsignificant genes are shown in gray. The horizontal dashed line indicates the significance threshold (adjusted p = 0.05). Selected genes associated with inflammatory responses and renal tubular biology are labeled. **(C)** Gene Ontology (GO) biological process enrichment analysis of genes exhibiting lower expression in Tolvaptan-treated kidneys compared with Placebo-treated kidneys 7 days after IRI. Enrichment analysis was performed using Enrichr (GO Biological Process 2025) on 164 genes identified by DESeq2 (adjusted p < 0.05). Bars represent -log10(adjusted enrichment p-value) for selected significantly enriched biological processes (adjusted p < 0.05). Numbers beside the bars indicate the number of contributing genes for each GO term. **(D)** Heatmap showing expression of selected inflammation-and immune response-associated genes in contralateral (C) and injured (I) kidneys from Placebo-and Tolvaptan-treated mice at day 7 after IRI. Log-normalized transcripts per million (TPM) values were scaled from 0 to 1 for visualization. **(E, F)** qRT-PCR analysis of selected chemokine and immune cell-associated genes in contralateral and injured kidneys from Placebo-and Tolvaptan-treated mice at day 7 after IRI. Gene expression is shown as 2^−ΔΔCt^. Statistical significance was assessed by two-way ANOVA followed by Tukey’s multiple-comparisons test. **(G)** Representative immunofluorescence images and quantification of F4/80-positive macrophage infiltration in the renal medulla of injured kidneys from Placebo-and Tolvaptan-treated mice at day 7 after IRI. F4/80 is shown in white and nuclei in blue. Scale bar, 20 µm. Data are shown as mean ± SEM, unpaired two-tailed Welch’s *t*-test. Each dot in E-G represents an individual mouse. *P < 0.05; **P < 0.01; ***P < 0.001; ns, not significant.

Bulk RNA sequencing of injured kidneys identified 307 differentially expressed genes between Tolvaptan-and Placebo-treated mice (adjusted p < 0.05), of which 164 exhibited lower and 143 higher expression in Tolvaptan-treated kidneys (Figure 4B, Supplementary Table 3S). Gene Ontology (GO) biological process enrichment analysis of genes with lower expression in Tolvaptan-treated kidneys revealed significant enrichment of immune-and inflammation-associated processes, including inflammatory response, positive regulation of cytokine production, and interferon-associated responses (Figure 4C, Supplementary table S4). Consistent with these transcriptional findings, several genes associated with monocyte recruitment and macrophage activation^21^ were strongly induced following injury but exhibited reduced expression in Tolvaptan-treated mice compared with placebo-treated controls (Figure 4D).

Targeted quantitative RT-PCR confirmed significantly lower expression of key chemokines and chemokine receptors involved in monocyte recruitment, including *Ccl2*, *Ccr2*, *Cx3cr1*, and *Cx3cl1*, in kidneys from Tolvaptan-treated mice following IRI (Figure 4E). In parallel, expression of macrophage-associated genes such as *Adgre1 (F4/80)*, *Cd68*, *Cd14*, and *Cd44* was also reduced compared with placebo-treated controls (Figure 4F).

To determine whether these transcriptional changes were reflected at the tissue level, macrophage accumulation was assessed by immunofluorescence staining for F4/80.

Tolvaptan-treated mice exhibited a significant reduction in F4/80-positive area within the renal medulla compared with control mice after IRI (Figure 4G). Together, these data indicate that pharmacological reduction of medullary osmolality induced by Tolvaptan is associated with attenuation of inflammatory cell recruitment and activation during the early recovery phase following ischemic kidney injury.

### Attenuation of inflammatory responses is associated with reduced fibrotic remodeling after ischemia-reperfusion injury

Persistent inflammation during the recovery phase of acute kidney injury is a key driver of maladaptive repair and fibrotic remodeling^2^. We therefore assessed whether the reduced inflammatory response observed in Tolvaptan-treated mice following IRI was associated with altered fibrotic outcomes at later stages.

Bulk RNA sequencing of injured kidneys at 7 days post-injury revealed reduced expression of genes associated with profibrotic signaling, extracellular matrix organization, and tissue remodeling in Tolvaptan-treated mice compared with placebo-treated controls. This included canonical fibrosis-associated genes, such as *Tgfb1*, *Col1a1*, *Fn1*, and *Vim*, together with a broader panel of extracellular matrix and matrix-remodeling genes (Figure 5A). Consistent with these transcriptomic findings, quantitative RT-PCR confirmed lower expression of selected fibrosis-associated genes in Tolvaptan-treated injured kidneys (Figure 5B).

**Figure 5.**
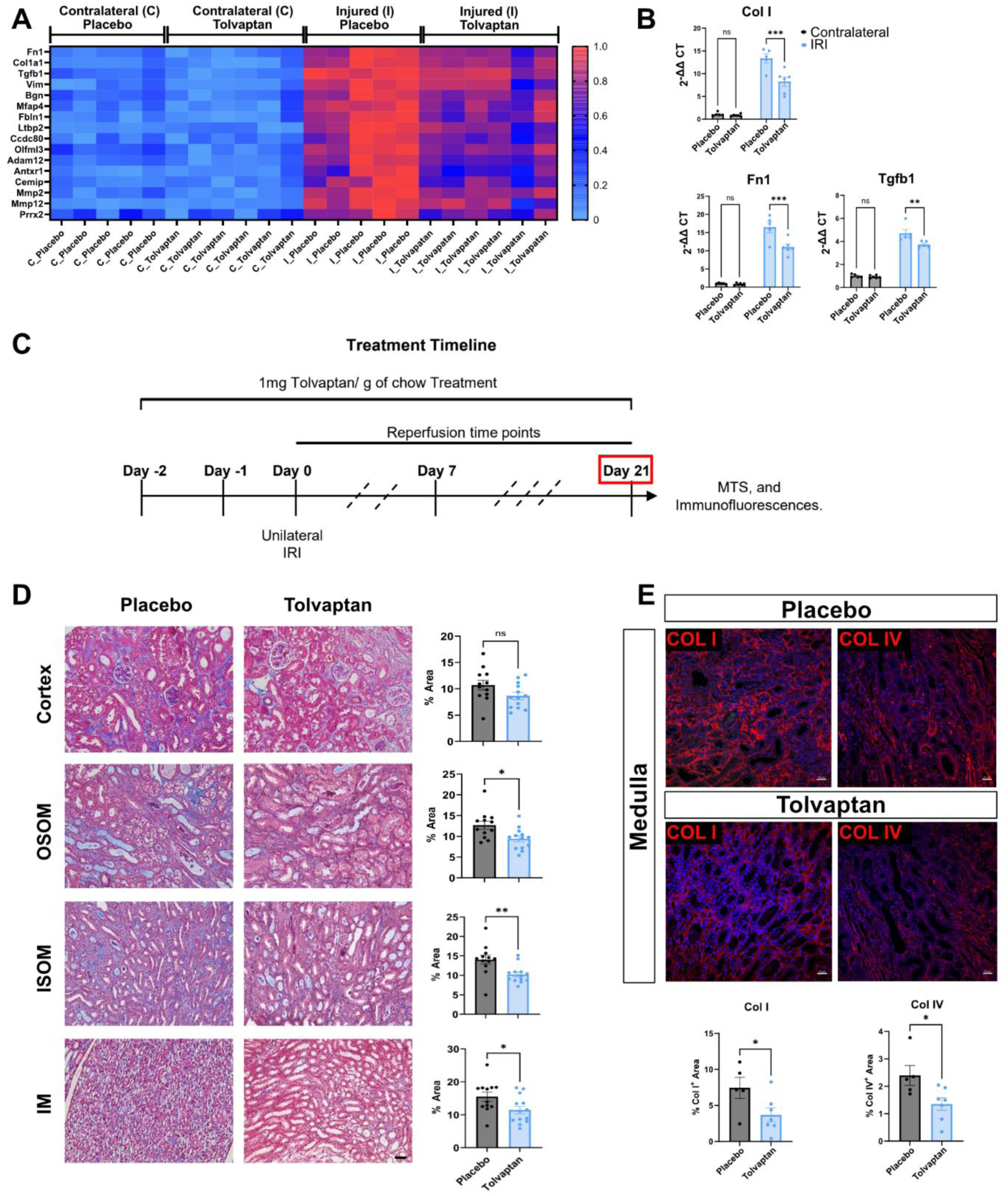
Effects of Tolvaptan treatment on renal fibrosis following ischemia– reperfusion injury. **(A)** Heatmap showing expression of fibrosis-associated genes in contralateral (C) and injured (I) kidneys from Placebo-(control diet without Tolvaptan) and Tolvaptan-treated mice at day 7 after IRI. Log-normalized transcripts per million (TPM) values were scaled from 0 to 1 for visualization. **(B)** qRT-PCR analysis of selected fibrosis-associated genes in contralateral and injured kidneys from Placebo-and Tolvaptan-treated mice at day 7 after IRI. Gene expression is shown as 2^−ΔΔCt^. **(C)** Experimental design for assessment of renal fibrosis at day 21 after unilateral IRI. Adult male C57BL/6N mice received Tolvaptan (1 mg/g chow) or Placebo starting 48 hours before IRI and continuing until tissue collection. **(D)** Representative Masson’s trichrome staining across injured kidneys and quantification of fibrotic area from Placebo-and Tolvaptan-treated mice at day 21 after IRI. Scale bars, 20 µm. **(E)** Representative immunofluorescence images and quantification of collagen I and collagen IV deposition in the renal medulla of injured kidneys from Placebo-and Tolvaptan-treated mice at day 21 after IRI. Scale bars, 20 µm. Each dot in B, D, E represents an individual mouse. Data are shown as mean ± SEM. Statistical significance in B was assessed by two-way ANOVA followed by Tukey’s multiple-comparisons test; statistical significance in D, E was assessed using an unpaired two-tailed *t*-test. \**P* < 0.05; **P < 0.01; ns, not significant.

To determine whether these transcriptional differences translated into altered tissue remodeling, kidneys were examined at 21 days post-IRI (Figure 5C), a time point corresponding to the transition from repair to chronic fibrotic remodeling. Histological assessment using Masson’s trichrome staining demonstrated reduced interstitial fibrosis in the renal medulla of Tolvaptan-treated mice compared with placebo-treated controls (Figure 5D). Consistently, immunofluorescence analysis revealed decreased deposition of Collagen I and Collagen IV within the medullary interstitium of Tolvaptan-treated kidneys (Figure 5E).

These findings indicate that modulation of medullary osmolality by Tolvaptan treatment during the injury and early repair phases is associated with sustained attenuation of fibrotic remodeling in the renal medulla following ischemia-reperfusion injury.

### Tolvaptan attenuates tubular injury, inflammatory activation and early fibrosis in cisplatin-induced nephropathy

We next evaluated the impact of Tolvaptan treatment in a mechanistically distinct kidney injury model induced by repeated low-dose cisplatin administration. Mice received three weekly injections of cisplatin (9 mg/kg), and kidneys were harvested three days after the final dose (Figure 6A). Analyses of kidney function parameters serum creatinine and urea (Figure 6B), kidney epithelial injury/failed repair markers *Havcr1* and *Vcam1* mRNA (Figure 6C) and fibrosis marker *Col1a1* mRNA (Figure 6D) indicated a suppression of cisplatin-induced kidney injury and early fibrosis by Tolvaptan. Consistently, immunofluorescence analysis of the outer stripe of the outer medulla demonstrated a marked induction of the tubular injury marker KIM-1 (*Havcr1*) in cisplatin-treated control mice, which was nearly abolished in Tolvaptan-treated animals (Figure 6E). In parallel, macrophage accumulation assessed by F4/80 staining was significantly reduced in the outer stripe of the outer medulla of Tolvaptan-treated mice compared with cisplatin-treated placebo-controls (Figure 6E). To address the possibility that Tolvaptan might interfere with cisplatin’s biological activity, we assessed cisplatin-induced cytotoxicity in the cervical cancer cell lines CaSki and SiHa, and in the ovarian cancer cell line OVCAR3 cells in the presence of Tolvaptan (1 and 3 µM). Across orthogonal readouts (live-cell imaging, Annexin V assay, and flow cytometry-based quantification of surviving mCherry-positive cells), Tolvaptan did not modify cisplatin-induced cell death (Supplementary Fig. S3). Together, these data strongly suggest that tubular injury, inflammatory activation and early fibrosis development during cisplatin-induced nephrotoxic injury are attenuated by Tolvaptan via a reduction of vasopressin-dependent kidney medullary osmolality.

**Figure 6.**
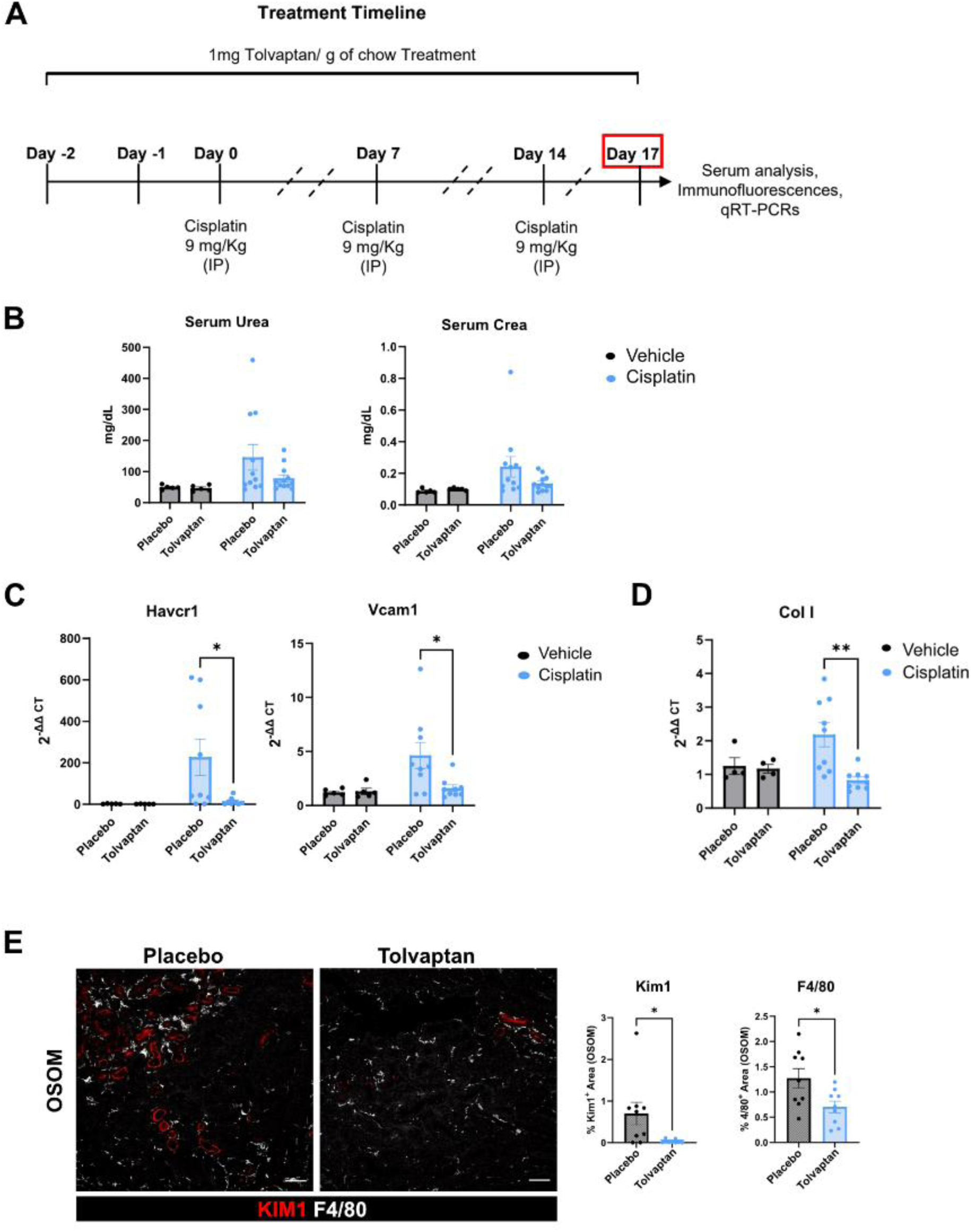
Effects of Tolvaptan treatment in a repeated low-dose cisplatin model of kidney injury. **(A)** Experimental design. Adult male C57BL/6N mice received Tolvaptan (1 mg/g chow) or Placebo (control diet without Tolvaptan) starting 48 hours before the first cisplatin administration. Cisplatin (9 mg/kg) was administered once weekly for three consecutive weeks, followed by sample collection at the indicated endpoint. **(B)** Serum creatinine and urea concentrations in Placebo-and Tolvaptan-treated mice following repeated cisplatin administration. **(C, D)** qRT-PCR analysis of selected kidney injury-, inflammation-, and fibrosis-associated genes in kidneys from Placebo-and Tolvaptan-treated mice. Gene expression is shown as 2^−ΔΔCt^. **(E)** Representative immunofluorescence images and quantification of KIM-1-positive tubular injury and F4/80-positive macrophage infiltration in the outer stripe of the outer medulla (OSOM) of Placebo-and Tolvaptan-treated mice. Scale bars, 20 µm. Each dot in B-E represents an individual mouse. Data are shown as mean ± SEM. Statistical significance in B-D was assessed by two-way ANOVA followed by Tukey’s multiple-comparisons test. Statistical significance in E was assessed using an unpaired two-tailed Welch’s *t*-test. \**P* < 0.05; **P < 0.01; ns, not significant.

## DISCUSSION

In this study, we demonstrate that vasopressin-dependent renal medullary hyperosmolarity is closely linked to tubular stress responses, inflammatory activation, and fibrotic remodeling after kidney injury. By pharmacologically reducing medullary osmolality using Avpr2 antagonism, we show that the medullary injury environment can be modulated in a manner that influences early injury trajectories and downstream repair outcomes.

A central observation of our work is that Tolvaptan lowers renal medullary osmolality and attenuates stress-associated transcriptional responses in medullary collecting duct cells. These changes extend beyond canonical vasopressin-responsive genes such as Aqp2 and include Nupr1, Ddit3, and Hmgb1. Previous studies have demonstrated that hyperosmolality induces an unfolded protein response in collecting duct principal cells^22^. Furthermore, CHOP, encoded by Ddit3, has been implicated in renal IRI, with prominent induction in the outer medulla and reduced injury following global CHOP deletion^23^. Similarly, tubular epithelial Hmgb1 deletion has been shown to improve postischemic recovery and attenuate AKI-to-CKD transition^24^. Although these findings support the biological relevance of the transcriptional changes observed in our study, they do not establish that reduced expression of these genes mediates the protective effects of Tolvaptan. Rather, our results indicate that AVPR2 antagonism is associated with altered epithelial stress responses in the setting of reduced medullary osmolality.

Following ischemia-reperfusion injury, reduction of medullary osmolality was associated with attenuated inflammatory signaling and reduced macrophage accumulation in the renal medulla. Inflammatory monocyte recruitment and macrophage activation are well-recognized contributors to maladaptive repair and fibrosis after AKI^25–27^. Our data indicate that modulation of the medullary osmotic *milieu* influences these processes during the early recovery phase, preceding overt fibrotic remodeling.

Consistent with this interpretation, Tolvaptan-treated mice exhibited reduced fibrotic remodeling of the renal medulla at later stages following IRI. While fibrosis is a multifactorial process, the sustained attenuation of extracellular matrix deposition observed here suggests that early modulation of injury-associated inflammation can have durable consequences for tissue remodeling. Notably, these effects were most prominent in the medulla, underscoring the importance of regional physiological context.

To assess whether these findings extend beyond ischemic injury, we examined a cisplatin-induced nephropathy model. Cisplatin injury is mechanistically distinct from IRI^28^ and is characterized by heterogeneous injury severity and variable functional readouts. In this setting, Tolvaptan reduced tubular injury markers and inflammatory activation in the outer stripe of the outer medulla, even when early changes in serum creatinine and urea were variable. These observations support the concept that medullary osmolality modulates intrinsic injury responses across different forms of AKI.

Our data support a model in which the medullary osmotic environment acts as a contextual modifier of injury-associated stress and inflammation. Further work will be required to define the precise signaling mechanisms involved and to determine how these processes interact with sex, chronic injury, and long-term functional outcomes.

Our study has important limitations. First, all experiments were performed in male mice, limiting the generalizability of our findings across sexes. Second, Tolvaptan was administered prophylactically, beginning before kidney injury. Therefore, our data do not establish whether treatment initiated after injury onset would have therapeutic efficacy. Finally, Tolvaptan was the sole intervention used to modify the renal medullary osmotic gradient. Consequently, we cannot fully distinguish effects attributable to reduced medullary osmolality from other consequences of AVPR2 antagonism.

In conclusion, our findings highlight vasopressin-dependent renal medullary hyperosmolality as a modifiable component of the kidney injury microenvironment. Tolvaptan treatment was associated with reduced tubular stress, inflammatory activation, and fibrotic remodeling after AKI. These results underscore the importance of considering regional kidney physiology when investigating mechanisms of injury and repair.

## AUTHOR CONTRIBUTIONS

KILC, FJB, and KMSO designed the study; KILC, FJB, JL, SC, CH, LN conducted experiments; KILC, FJB, SC, CH, LN, analyzed data; KILC, AS, KMSO provided resources; KILC, FJB, LN made the figures; KILC wrote the manuscript; KMSO reviewed and edited the manuscript; all authors reviewed and approved the final version of the manuscript.

## FUNDING

This project was supported by grants awarded to KMSO: German Research Foundation (DFG) Research Training Group GRK 2318 (ID 318905415); DFG Research Unit FOR 2841 (ID 400728090); DFG Collaborative Research Grant SFB 1365 (ID 394046635), DFG research grant (ID 547091997), and grants awarded to KILC: internal funding (HiLF) of the MHH, and Young Investigator Research Funding (FOR2841, DFG).

## ACKNOWLEDGMENTS

We thank the Animal Phenotyping Platform at the MDC for their excellent assistance. We extend our gratitude to Tatjana Luganskaja (MDC), Gabriele N’Diaye (MDC), and Martina Flechsig (MHH) for their excellent technical support. We thank Dr. Michael Morgan and Dr. Ivana Kutle (MHH) for generating the mCherry-expressing cancer cell lines used in this study.

## DATA AVAILABILITY

Processed single-nucleus RNA sequencing data and an interactive Shiny application for gene-level exploration of expression changes across all annotated renal cell populations are available at http://shiny.mdc-berlin.de/Tol/, login data: Username: Schmidt-Ott, Password: Tolvaptan. Raw and processed single-nuclei and bulk sequencing data have been deposited to Gene Expression Omnibus (GEO), accession numbers: GSE350654 and GSE350655. All additional raw data supporting the findings of this study, together with the code used for analysis, have been deposited in the MHH Research Data Repository (https://doi.org/10.26068/mhhrpm/20261021-000) and are available upon request. No new codes were developed specifically for this study.

## Notes

### Competing Interest Statement

The authors have declared no competing interest.

