## Supplemental information for "Targeting the Kidney’s Osmotic Gradient with AVPR2 Antagonism Limits Inflammation and Fibrosis after Injury"

**Table of contents of supplemental material:**

• Supplementary Figure S1. Quality control metrics for single-nucleus RNA sequencing datasets.
• Supplementary Figure S2. Stress-associated genes responsive to Tolvaptan in mouse kidney are increased in human AKI.
• Supplementary Figure S3. Tolvaptan does not attenuate cisplatin-induced tumor cell death *in vitro*.
• Supplementary Table S1. Antibodies used for immunofluorescence.
• Supplementary Table S2. Primers used for qRT-PCR.
• Supplementary Table S3. DESeq2 differential gene expression analysis of kidneys from tolvaptan- and placebo-treated mice 7 days after ischemia–reperfusion injury.
• Supplementary Table S4. Gene Ontology biological process enrichment analysis of genes downregulated in kidneys from tolvaptan-treated mice 7 days after ischemia–reperfusion injury.

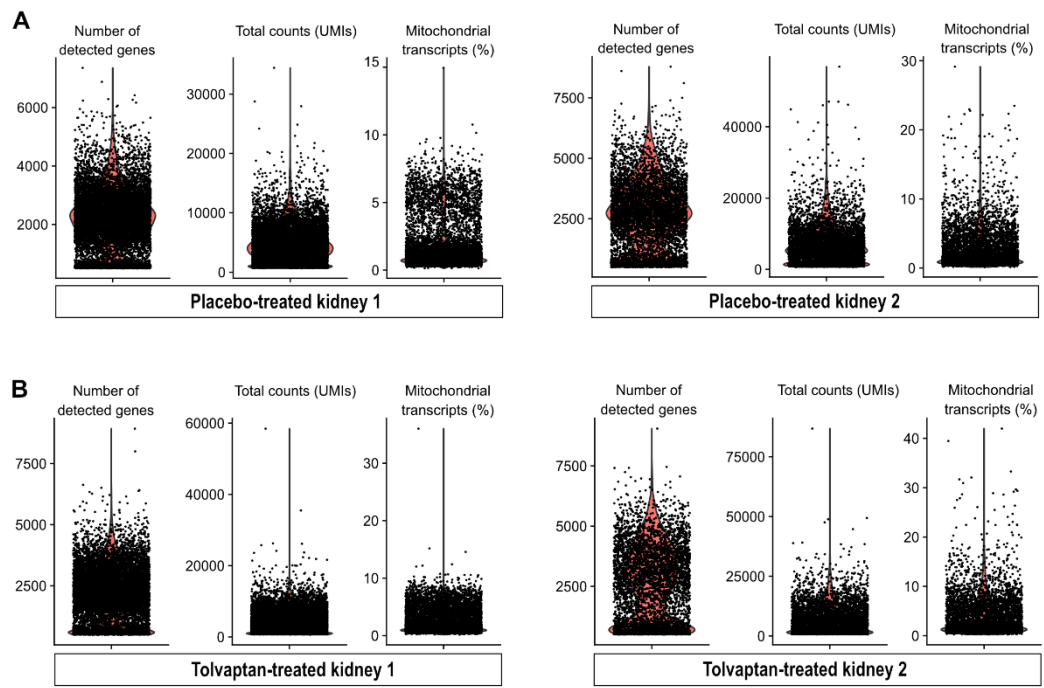

**Supplementary Figure S1. Quality control metrics for single-nucleus RNA sequencing datasets.** (A) Distribution of the number of detected genes (nFeature\_RNA), total transcript counts (nCount\_RNA; UMIs), and percentage of mitochondrial transcripts across nuclei from kidneys from placebo-treated mice (kidney 1 and 2). (B) Corresponding quality control metrics from kidneys from tolvaptan-treated mice (kidney 1 and 2). Each point represents an individual nucleus; violin plots indicate the distribution density.

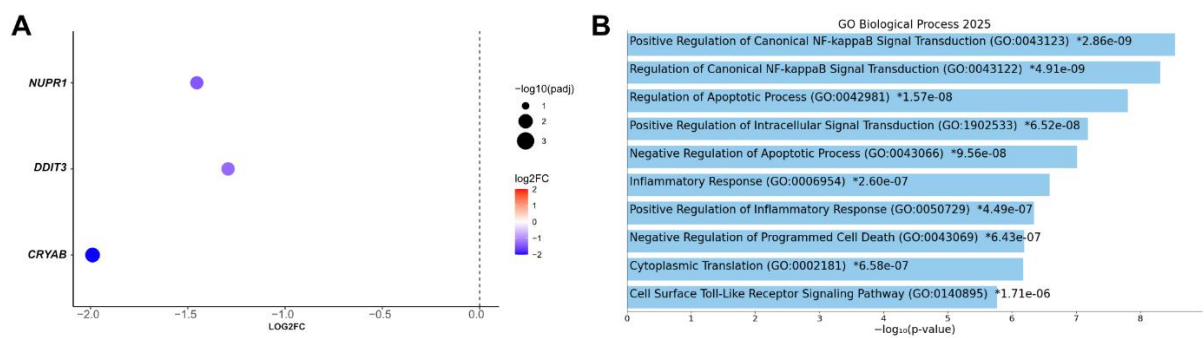

**Supplementary Figure S2. Stress-associated genes responsive to Tolvaptan in mouse kidney are increased in human AKI.** Analysis of publicly available human kidney transcriptomic datasets<sup>1</sup> was performed to assess gene expression changes in acute kidney injury (AKI). Control samples included tumor-adjacent normal kidney tissues from nephrectomies and post-mortem biopsies from a non-AKI individual (15-120 min after cessation of circulation). AKI samples consisted of post-mortem kidney biopsies (60-120 min) from individuals with severe AKI (KDIGO stage 2-3; AKI1-AKI8). (A) Expression of stress-associated genes, including *CRYAB*, *DDIT3*, and *NUPR1*, is increased in AKI compared with

controls. Fold changes are shown as $\log_2(\text{control versus AKI})$, where negative values indicate higher expression in AKI. **(B)** Gene Ontology (GO) biological process enrichment analysis of genes upregulated in AKI (adjusted $p < 0.05$ and fold change $\leq 0.5$) shows enrichment of inflammatory signaling, apoptotic processes, and stress-response pathways, including canonical NF-κB signaling. Bars represent $-\log_{10}(p\text{-value})$ for the top enriched terms.

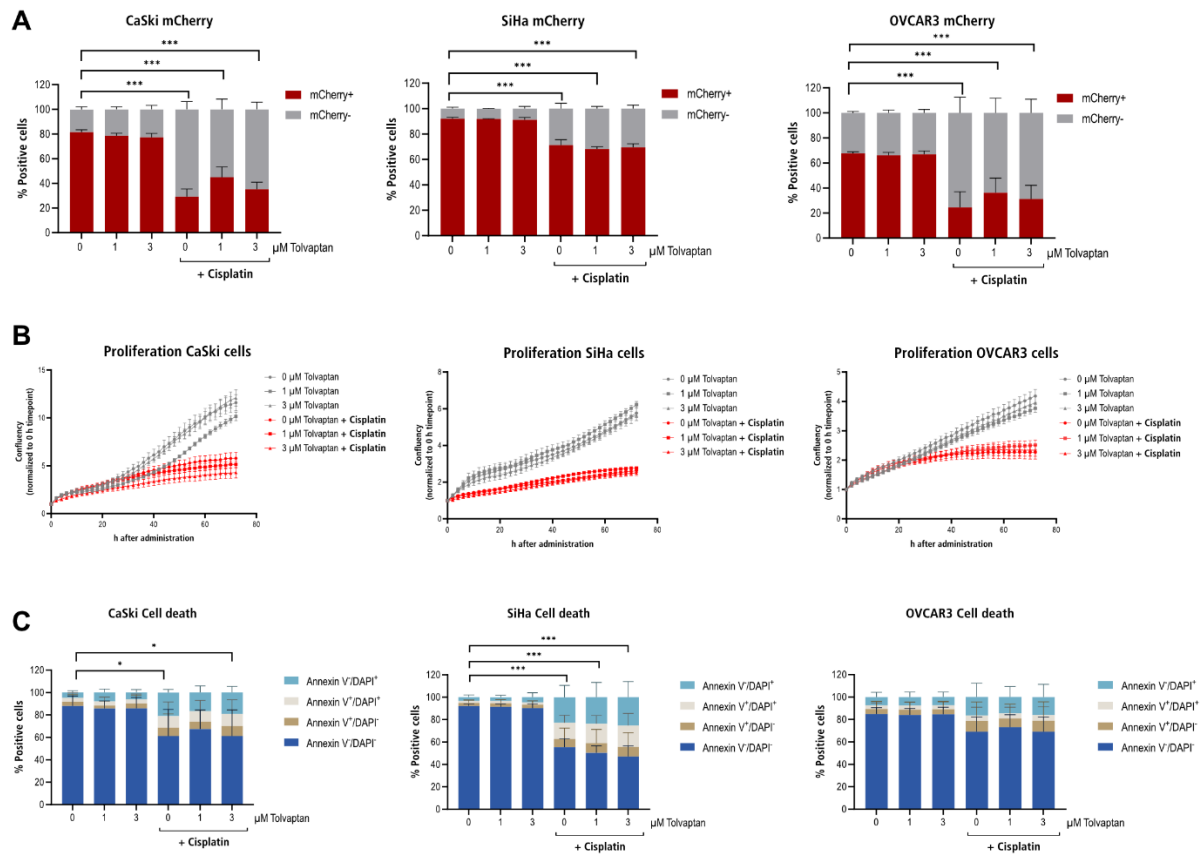

**Supplementary Figure S3. Tolvaptan does not attenuate cisplatin-induced tumor cell death *in vitro*.** **(A-C)** mCherry-labeled human cervical carcinoma (CaSki and SiHa) and ovarian carcinoma (OVCAR3) cells were treated with cisplatin (10 μM) in the presence or absence of Tolvaptan (1 or 3 μM) for 72 hours. **(A)** Flow cytometric quantification of mCherry-positive cells, expressed as the percentage of mCherry-positive cells following treatment. **(B)** Live-cell imaging-based analysis of cell proliferation during the 72-hour treatment period. **(C)** Flow cytometric analysis of Annexin V-positive cells, expressed as the percentage of positive cells following treatment. Data are presented as mean ± SD. Statistical significance was assessed by two-way ANOVA followed by Dunnett's multiple-comparisons test. \*$P < 0.05$; \*\*\*$P < 0.001$.

**Supplementary Table S1.** Antibodies used for immunofluorescence.

| Target Protein | Host | Type | Dilution | Supplier | Catalog No. |
| --- | --- | --- | --- | --- | --- |
| **Collagen I** | Rabbit | Primary | 1:300 | Meridian Life Science | T40777R |
| **Collagen IV** | Goat | Primary | 1:200 | SouthernBiotech | 1340-01 |
| **F4/80** | Rabbit | Primary | 1:300 | Cell Signaling | 70076T |
| **Kim1** | Goat | Primary | 1:200 | R&D Systems | AF1817 |
| **Alexa Fluor™ 647 Donkey anti-Rabbit IgG (H+L)** | Donkey | Secondary | 1:1000 | Invitrogen | A-31573 |
| **Cy™3 AffiniPure Donkey Anti-Goat IgG (H+L)** | Donkey | Secondary | 1:1000 | Jackson ImmunoResearch | 705-165-147 |

**Supplementary Table S2.** Primers used for qRT-PCR.

| Target Gene | Forward (5'-3') | Reverse (5'-3') |
| --- | --- | --- |
| *Hprt* | CAGTCCCAGCGTCGTGATTA | AGCAAGTCTTTCAGTCCTGTC |
| *Adgre1* | AAACTCAAGGACACGAGGTTGC | ACAGGTATGCCATGATGCTTGC |
| *Ccl2* | TTAAAAACCTGGATCGGAACCAA | GCATTAGCTTCAGATTTACGGGT |
| *Ccr2* | ATCCACGGCATACTATCAACATC | TCGTAGTCATACGGTGTGGTG |
| *Cx3cl1* | ACGAAATGCGAAATCATGTGC | CTGTGTCGTCTCCAGGACAA |
| *Cx3cr1* | GAGTATGACGATTCTGCTGAGG | CAGACCGAACGTGAAGACGAG |
| *Cd68* | ACCTACATCAGAGCCCGAGTACAG | TCTGCGCCATGAATGTCCACTG |
| *Cd14* | TGAATTGGGCGAGAGAGGACTG | AACTTGGAGGGTCGGGAACTTG |
| *Cd44* | ACACCTACCTTCCTACTGAACAGC | TGGATGAGTCTCGATCTCTGGTG |
| *Vcam1* | TTCGGTTGTTCTGACGTGTG | TACCACCCCATTGAGGGGAC |
| *Col1a1* | CCG CTG GTC AAG ATG GTC | CCT CGC TCT CCA GCC TTT |
| *Tgfb1* | TTG CTT CAG CTC CAC AGA GA | TGG TTG TAG AGG GCA AGG AC |
| *Fn1* | GAT ACA TGA CCC CTT CAT TG | CGG TAG GAC CTT CTA TTC CT |
| *Havcr1* | AAACCAGAGATTCCCACACG | GTCGTGGGTCTTCCTGTAGC |

**Supplementary Table S3.** DESeq2 differential gene expression analysis of kidneys from tolvaptan- and placebo-treated mice 7 days after ischemia–reperfusion injury.

Log2 fold changes represent tolvaptan relative to placebo, with positive values indicating higher expression in tolvaptan-treated mice. Adjusted P values were calculated using the Benjamini–Hochberg method. This table is provided as an Excel file.

**Supplementary Table S4.** Gene Ontology biological process enrichment analysis of genes downregulated in kidneys from tolvaptan-treated mice 7 days after ischemia–reperfusion injury.

Gene Ontology (GO) biological process enrichment analysis was performed using Enrichr (GO Biological Process 2025) on 164 genes showing significantly lower expression in tolvaptan-treated mice compared with placebo-treated mice (DESeq2 adjusted $P < 0.05$). A background of 14,997 genes was used. The table presents the enriched GO terms and their associated enrichment statistics. This table is provided as an Excel file.
